# Reduced SARS-CoV-2 mRNA vaccine immunogenicity and protection in mice with diet-induced obesity and insulin resistance

**DOI:** 10.1101/2022.12.07.519460

**Authors:** Timothy R. O’Meara, Etsuro Nanishi, Marisa E. McGrath, Soumik Barman, Danica Dong, Carly Dillen, Manisha Menon, Hyuk-Soo Seo, Sirano Dhe-Paganon, Robert K. Ernst, Ofer Levy, Matthew B. Frieman, David J. Dowling

**Author notes:** **Corresponding author:** David J. Dowling, Precision Vaccines Program, Division of Infectious Diseases, Boston Children’s Hospital; Harvard Medical School, Rm 842, Boston, MA 02115, USA. These authors contributed equally to this manuscript (Timothy R. O’Meara, Etsuro Nanishi, Marisa E. McGrath). Co-senior authors (Ofer Levy, Matthew B. Frieman, David J. Dowling). **Author contributions:** T.R.O. and E.N. conceived, designed, performed, and analyzed the experiments and wrote the manuscript; M.E.M. designed, performed, and analyzed SARS-CoV-2 neutralization experiments and mouse challenge study and wrote the paper; S.B. and M.M. performed splenocyte restimulation; S.B. performed flow cytometry and the analysis; D.D. performed qPCR experiments and their analysis; C.D., R.M.J., and M.B.F. performed and analyzed SARS-CoV-2 neutralization experiments and mouse challenge study; H.S. and S.D.P. produced the RBD protein; R.K.E. contributed to the design of experiments; O.L., M.B.F., and D.J.D. conceived the project, designed the experiments, assisted in interpretation of the results, and edited the manuscript.

## Abstract

**Background:** Obesity and Type 2 Diabetes Mellitus (T2DM) are associated with an increased risk of severe outcomes from infectious diseases, including COVID-19. These conditions are also associated with distinct responses to immunization, including an impaired response to widely used SARS-CoV-2 mRNA vaccines.

**Objective:** To establish a connection between reduced immunization efficacy via modeling the effects of metabolic diseases on vaccine immunogenicity that is essential for the development of more effective vaccines for this distinct vulnerable population.

**Methods:** We utilized a murine model of diet-induced obesity and insulin resistance to model the effects of comorbid T2DM and obesity on vaccine immunogenicity and protection.

**Results:** Mice fed a high-fat diet (HFD) developed obesity, hyperinsulinemia, and glucose intolerance. Relative to mice fed a normal diet (ND), HFD mice vaccinated with a SARS-CoV-2 mRNA vaccine exhibited significantly lower anti-spike IgG titers, predominantly in the IgG2c subclass, associated with a lower type 1 response, along with a 3.83-fold decrease in neutralizing titers. Furthermore, enhanced vaccine-induced spike-specific CD8^+^ T cell activation and protection from lung infection against SARS-CoV-2 challenge were seen only in ND mice but not in HFD mice.

**Conclusion:** We demonstrate impaired immunity following SARS-CoV-2 mRNA immunization in a murine model of comorbid T2DM and obesity, supporting the need for further research into the basis for impaired anti-SARS-CoV-2 immunity in T2DM and investigation of novel approaches to enhance vaccine immunogenicity among those with metabolic diseases.

**Capsule summary:** Obesity and type 2 diabetes impair SARS-CoV-2 mRNA vaccine efficacy in a murine model.

## INTRODUCTION

The size and proportion of the population with obesity and diabetes mellitus (DM) are growing across the globe, especially in high-income countries. Among US adults, the prevalence of obesity and DM are 41.9% and 14.8%, respectively^1^. The relationship between DM and an increased risk of morbidity and mortality caused by a variety of infectious diseases has long been recognized, especially in older adults with DM^2^. Similarly, DM and obesity are risk factors of severe COVID-19 or death, along with other factors such as older age, male sex, and underlying comorbidities (e.g., cardiovascular disease and chronic kidney disease)^3-6^. The prevalence of these metabolic disorders indicates an urgent need to prevent the incidence of severe infections, specifically COVID-19, in these vulnerable populations to reduce disease burden.

Despite improving overall disease outcomes, many currently approved vaccines, including the SARS-CoV-2 BNT162b2 mRNA vaccine, are not as effective in patients with DM or obesity. Following the introduction of mRNA vaccines against SARS-CoV-2, clinical studies found that Type 2 DM (T2DM) is associated with significant reductions in both humoral and cellular responses to vaccination against SARS-CoV-2, particularly among those with poor glycemic control^7, 8^. Reduced vaccine immunogenicity has been observed among adults, especially men, with obesity^9^. Together, these findings suggest that metabolic diseases impair vaccine responses and increase the risk of severe COVID-19. However, the exact effects of metabolic disease on the quality of humoral and cellular immune responses remain unclear. There is therefore a need to assess the causes of impaired vaccine response in those with metabolic disease and evaluate what aspects of immunity are affected to inform optimization of vaccine approaches for this vulnerable population.

While our understanding of the influence of obesity and T2DM on SARS-CoV-2 vaccine responses remains limited, murine models of diet-induced obesity (DIO) and insulin resistance have facilitated initial studies of the connections between metabolic disease, immunity, and viral disease pathology, particularly in the context of influenza. Following infection with influenza, DIO mice exhibited higher lung damage and mortality^10-14^. Further, DIO mice mounted impaired immune responses following immunization with subunit or inactivated-virus influenza vaccines, including decreased antibody (Ab) titers relative to controls, lower CD8^+^ T cell levels, impaired protection from live viral challenge, and greater waning in humoral immunity^15-18^. Additionally, studies of MERS-CoV and SARS-CoV-2 infection have found that DIO mice exhibit increased lung titers and/or greater morbidity and mortality following live-virus challenge relative to controls^19-21^. However, little is known regarding the effects of obesity and hyperglycemia on SARS-CoV-2 vaccine responses. Furthermore, no studies have yet evaluated the effects of obesity and hyperglycemia on mRNA vaccine immunogenicity in detail despite the widespread use of mRNA-based SARS-CoV-2 vaccines in the clinic.

We therefore sought to address these gaps by evaluating the effects of obesity and T2DM on Ab levels and function, T cell responses, and protection from live SARS-CoV-2 challenge in animals that received the SARS-CoV-2 BNT162b2 mRNA vaccine. To this end, we established a mouse model of obesity, hyperinsulinemia, and glucose intolerance using a high-fat diet (HFD) and then immunized mice with BNT162b2 or an alum adjuvanted SARS-CoV-2 spike receptor-binding domain (RBD) subunit vaccine. We then assessed binding and neutralizing Ab titers, CD8^+^ T cell activation, and protection from infection during viral challenge. We found that HFD-induced obesity and T2DM impaired both humoral and cellular immune responses post-BNT162b2 immunization. HFD-fed mice had significantly lower neutralizing titers and IgG2c titers compared to mice fed with a normal diet (ND). Furthermore, while ND mice exhibited RBD-specific CD8^+^ T cell activation, T cell activation profiles were not significantly enhanced in HFD mice when compared to a PBS-injected control. In line with these immunogenicity data, lung viral titers and inflammation profiles after viral challenge were only significantly reduced relative to the PBS-injected group among ND mice, while no significant reduction versus the PBS group was observed in the HFD group. Overall, our study demonstrates that diet-induced obesity and T2DM in a murine model reduce immunogenicity and protective efficacy of the SARS-CoV-2 BNT162b2 mRNA vaccine, laying the groundwork for further study of the mechanisms of these deficiencies and strategies that can be used to overcome them.

## RESULTS

### A high-fat diet causes weight gain, hyperinsulinemia, and glucose intolerance in male C57BL/6J mice

We first confirmed that feeding a HFD led to diet-induced obesity (DIO), fasting hyperinsulinemia, and glucose intolerance in male C57BL/6J mice as previously observed for this model^21-23^. To this end, mice were fed a HFD containing 60% kcal from fat or a ND containing 10% kcal from fat beginning at age 6 weeks (**table S1**). Animals were transferred from the supplier at 15 weeks old, allowed to acclimate for two weeks, and weighed weekly through the post-vaccination blood draw at 30 weeks of age. After feeding mice the HFD for 18 weeks, fasting serum insulin was measured. Glucose intolerance was measured via an intraperitoneal glucose tolerance test (IPGTT) the following week (**Fig. 1A**). As expected, mice that received a HFD were significantly heavier than mice fed a ND throughout the experiment (*P* < 0.0001 at all time points, **Fig. 1B**). The HFD mice were visually distinct from the ND mice, appearing much wider and rounder throughout the experiment (**fig. S1**). During the week of the prime vaccination, HFD mice ranged from 37.2 to 61.2 g, with an average weight of 48.8 g, while ND mice ranged from 26.9 to 36.8 g, averaging 31.8 g (**Fig. 1C**). In addition to weight gain, HFD mice also developed hyperinsulinemia, with significantly elevated fasting serum insulin levels as compared to ND mice at age 24 weeks (*P* < 0.0001, **Fig. 1D**). Further, serum insulin demonstrated a significant positive correlation with weight in both the ND mice (*r* = 0.4369, *P* = 0.0061) and HFD mice (*r* = 0.6194, *P* < 0.0001), suggesting an association between weight gain and hyperinsulinemia, particularly in mice that received a HFD (**Fig. 1E**). Finally, we employed an IPGTT to assess glucose intolerance in HFD mice versus ND mice (**Fig. 1F, G**). HFD mice had a significantly higher median blood glucose following a 6-hour fast than ND mice (244 mg/dL vs. 170 mg/dL, *P* < 0.0001, **Fig. 1G**). Following intraperitoneal injection of 2 g/kg dextrose, HFD mice maintained significantly higher blood glucose measurements than ND mice at 30, 75, and 120 minutes after injection (*P* < 0.0001 for all comparisons, **Fig. 1F**) and ended at a median of 600 mg/dL versus 225 mg/dL in ND mice (**Fig. 1G**). Moreover, 21 of the 38 HFD mice remained > 600 mg/dL, the blood glucose meter’s upper limit of detection, at the final time point (120 minutes after injection), in contrast with none of the ND mice (**Fig. 1G**).

**Figure 1.**
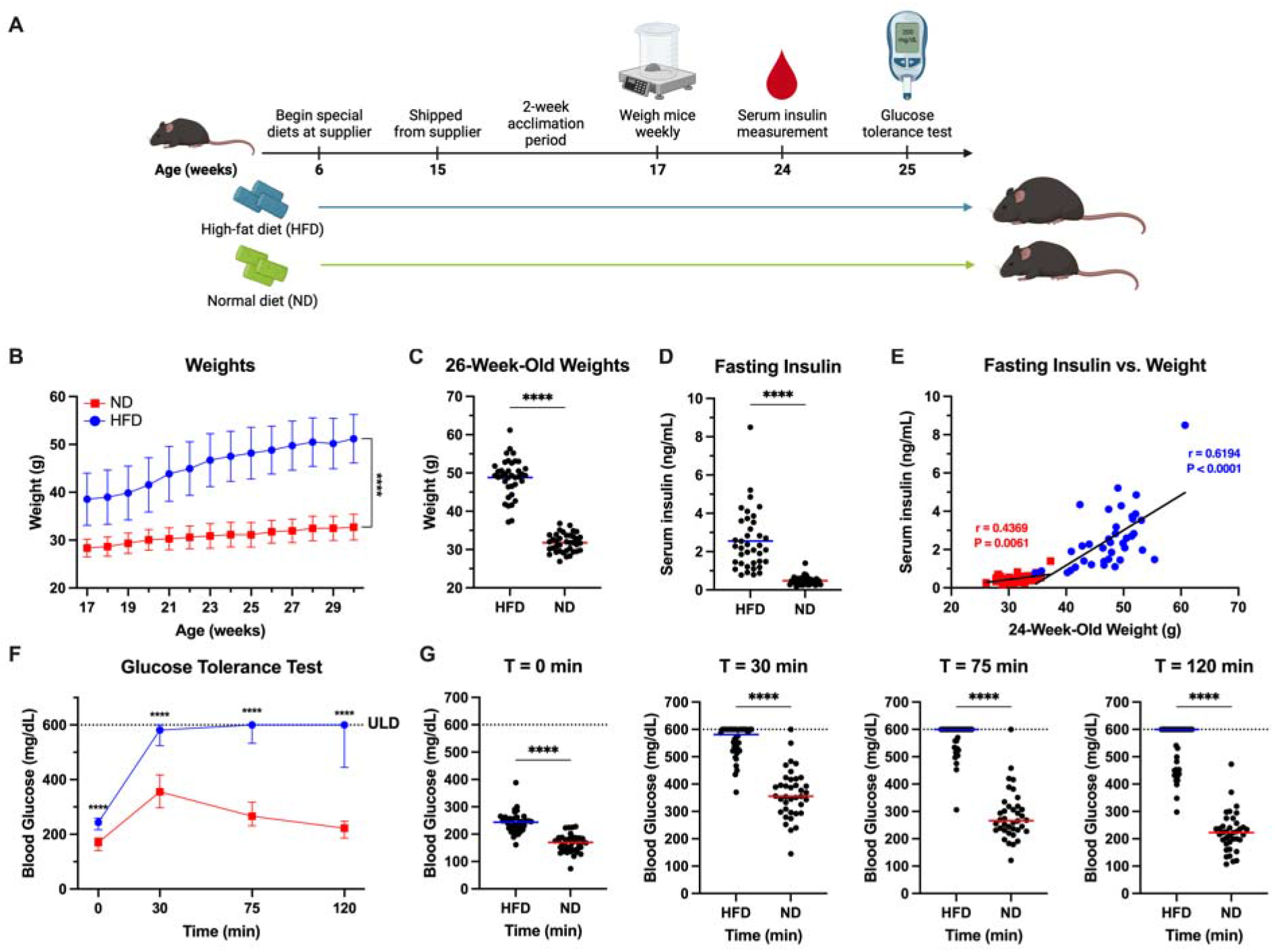
Male C57BL/6J mice fed a high-fat diet develop obesity, hyperinsulinemia, hyperglycemia, and poor glucose tolerance. Male C57BL/6J mice were fed a high-fat diet (HFD) consisting of 60% kcal from fat or an ingredient-matched control diet containing 10% kcal from fat (normal diet, ND) beginning at age 6 weeks. Mice were transferred from the vendor at 15 weeks old and allowed to acclimate for two weeks following receipt. Mice were then weighed weekly to assess weight gain. At 24 weeks old, serum insulin was measured by ELISA following a 6-hour fast. The following week, an intraperitoneal glucose tolerance test was performed to assess glucose tolerance. (**A**) Experimental design. (**B**) Mouse weights during the study. (**C**) Weights at 26 weeks of age. (**D**) Serum insulin levels measured at 24 weeks of age after a 6-hour fast. (**E**) Pearson’s correlation analysis was used to examine the correlation between serum fasting insulin levels and weights at 24 weeks of age. Lines indicate linear regression. (**F, G**) Glucose tolerance was assessed by measuring blood glucose at time points 0, 30, 75, and 120 min following a 6-hour fast and intraperitoneal injection of 2 g/kg dextrose. Blood glucose values above the glucometer’s 600 mg/dL upper limit of detection (ULD) were assigned a value of 600 mg/dL. N = 38 per group in all experiments. Longitudinal graphs display mean and standard deviation (**B)** or median and IQR (**F**). Bars represent means (**C, D**) or medians in all dot plots (**G**). Significance was assessed by unpaired t-test (**B–D**) or Mann-Whitney U-tests (**F, G**), correcting for multiple comparisons when relevant. **** *P* < 0.0001.

### HFD mice elicited impaired antibody responses following SARS-CoV-2 mRNA vaccination

Following establishment of the comorbid obesity, hyperinsulinemia, and glucose intolerance phenotypes, mice were immunized with SARS-CoV-2 BNT162b2 mRNA or a protein subunit vaccine benchmark vaccine at a 2-dose regimen with a 14-day interval to assess the effects of the HFD on vaccine immunogenicity and protective efficacy (**Fig. 2A**). Mice were randomly assigned to receive 1 µg of SARS-CoV-2 BNT162b2 mRNA (Comirnaty^®^), or 10 µg of recombinant monomeric SARS-CoV-2 spike RBD protein formulated with 100 µg of aluminum hydroxide (Alhydrogel^®^). Within the HFD or ND mice, each vaccine treatment group had generally comparable weights, insulin levels, and IPGTT results (**Fig. S2**). Immunizations were given intramuscularly to mice at 26 and 28 weeks of age. Two weeks after the 2^nd^ immunization, humoral immunity was assessed.

**Figure 2.**
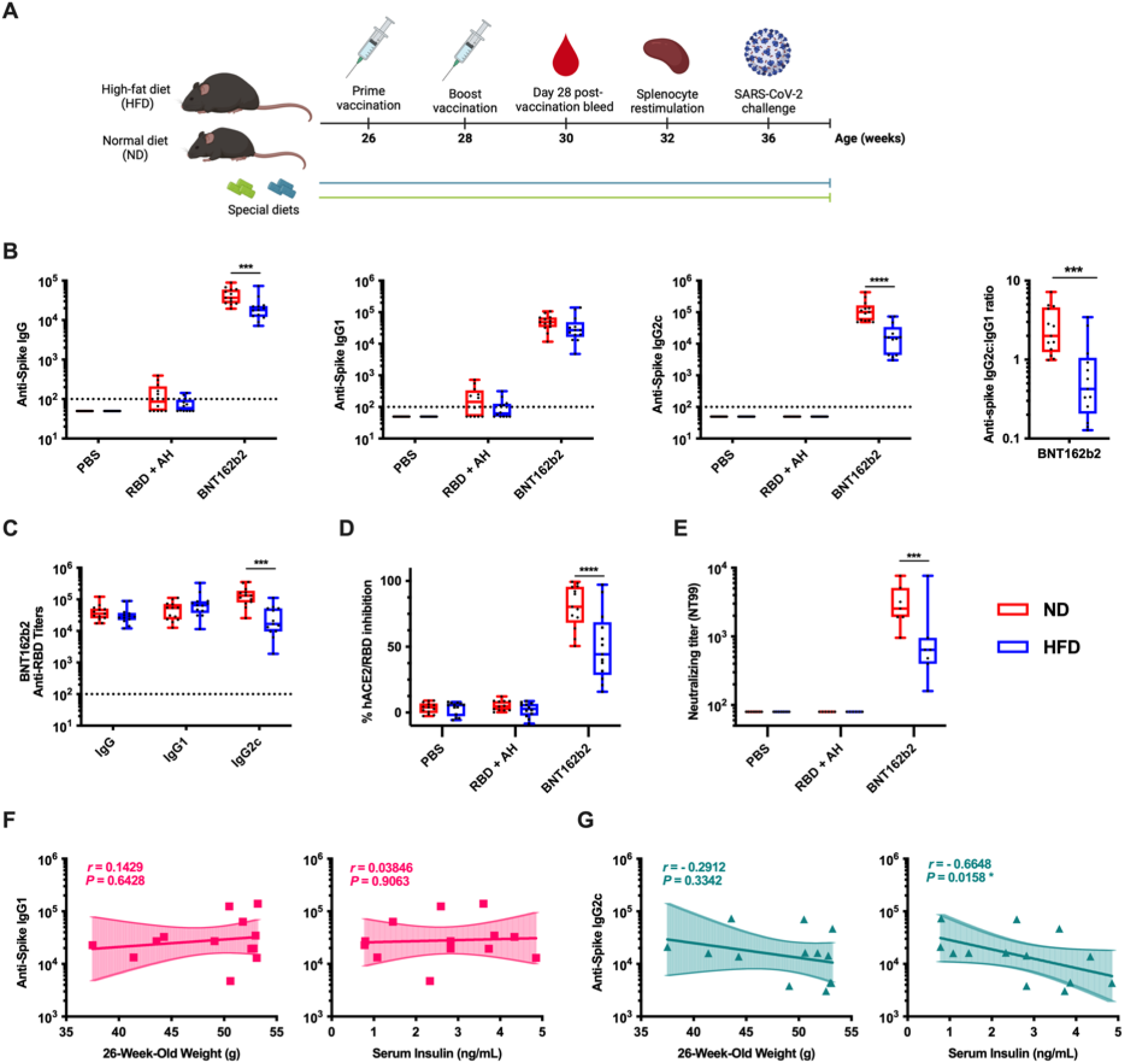
SARS-CoV-2 mRNA vaccine elicits reduced humoral immunogenicity in HFD mice. Male C57BL/6J mice fed with high-fat diet (HFD) or normal diet (ND) were immunized intramuscularly with a 2-dose regimen with 10 µg of aluminum-adjuvanted recombinant RBD or 1 µg of BNT162b2 mRNA. Serum samples were collected 14 days after the final immunization. **(A)** Experimental schematic. (**B–E**) Anti-Spike IgG, IgG1, IgG2c, and IgG2c:IgG1 ratio (**B**), Anti-RBD IgG, IgG1, and IgG2c post BNT162b2 mRNA immunization (**C**), hACE2-RBD inhibition rate (**D**), and WA1 SARS-CoV-2 neutralizing titers (**E**) were assessed. Dashed lines represent lower limits of detection. After log transformation, data were analyzed by two-way ANOVA followed by post-hoc tests for multiple comparisons. * *P* < 0.05, ** *P* < 0.01, *** *P* < 0.001, and **** *P* < 0.0001. (**F-G**) Correlations between anti-spike IgG1 (**F**) or IgG2c (**G**) titers and weight at vaccination or fasted serum insulin at 24 weeks of age of HFD mice following BNT162b2 mRNA immunization were assessed with Spearman’s rank correlation. Lines indicate linear regression. N = 12-13 per group except for neutralizing titers (N = 6-7 mice per group).

In ND mice, robust humoral responses were observed after BNT162b2 immunization, while an alum-adjuvanted RBD subunit vaccine induced limited Abs (**Fig. 2B–E**). Importantly, among mice that received BNT162b2, there was a 2.2-fold reduction in anti-spike IgG in HFD mice compared to ND mice (*P* = 0.0002, **Fig. 2B**). In further IgG subclass assessment, there was a large reduction in anti-spike IgG2c in HFD mice compared to ND mice post BNT162b2 immunization (GMTs of 105191 vs. 14250, *P* < 0.0001, **Fig. 2B**) although the difference in IgG1 titers was not significant. Accordingly, the Ab response was significantly skewed toward IgG2c in ND mice relative to HFD mice, with mean IgG2c:IgG1 ratios of 2.81 versus 0.87, respectively (*P* = 0.002, **Fig. 2B**). RBD plays a key role in ACE2 binding and is the main target of neutralizing Abs. We thus assessed anti-RBD IgG titers and confirmed a significant reduction in anti-RBD IgG2c in HFD mice compared to ND mice post BNT162b2 immunization (**Fig. 2C**). To analyze Ab function, we next performed a surrogate of virus neutralization test that measures the degree of inhibition of RBD binding to hACE2 by immune sera, as well as a neutralization assay with live SARS-CoV-2 virus. HFD mice exhibited significantly impaired hACE2-RBD binding inhibition and live virus neutralization relative to ND mice post BNT162b2 immunization (*P* < 0.0001 and *P* = 0.0003 respectively, **Fig. 2D, E**). To assess the effects of obesity and T2DM on humoral immunity individually, correlations between either weight at the time of immunization or fasting insulin, measures associated with obesity and diabetic phenotypes, respectively, and Ab responses were assessed in HFD mice. Interestingly, serum insulin, but not weight, negatively correlated with anti-spike IgG2c titers (*P* = 0.016 and *P* = 0.334 respectively, **Fig. 2F, G**). Overall, these results demonstrate that, relative to control mice fed a ND, HFD mice mount impaired Ab responses following immunization with BNT162b2, marked by a reduction in anti-spike IgG titers, a lower IgG2c:IgG1 ratio, impaired inhibition of hACE2-RBD binding, and a reduction in live virus neutralizing titers.

### HFD mice exhibit impaired CD8^+^ T cell activation following SARS-CoV-2 mRNA vaccination

SARS-CoV-2 spike specific CD8^+^ T cells are elicited by mRNA vaccines and contribute to protection against SARS-CoV-2 ^24-26^. We therefore analyzed spike RBD-specific CD8^+^ T cell responses of the HFD and ND mice 4 weeks after the final immunization. Splenocytes were collected and stimulated with overlapping peptides of the wildtype SARS-CoV-2 spike RBD. Intracellular expression of interferon-γ (IFNγ), TNF, and IL-2 in CD8^+^ T cells was assessed by flow cytometry to quantify antigen-specific cytotoxic T cell responses (**Fig. 3A**). As expected, BNT162b2 immunization elicited significantly higher CD8^+^ T cell expression of IFNγ, TNF, and IL-2 than PBS injection in ND mice (*P* = 0.002, *P* = 0.007, and *P* = 0.010, respectively), while alum-adjuvanted RBD subunit vaccine was not significant versus PBS for any cytokines (**Fig. 3B–D**). In contrast, neither BNT162b2 nor alum-adjuvanted RBD significantly induced CD8^+^ T cell IFNγ, TNF, or IL-2 expression versus PBS among HFD mice (**Fig. 3B–D**). While only the ND mice exhibited a significant vaccine-induced increase in cytokine expression over their corresponding PBS group, there was not a significant difference in cytokine expression comparing HFD and ND mice when compared head-to-head within the BNT162b2 vaccination condition (**Fig. 3B–D**). However, the median percentages of IFNγ, TNF, and IL-2 positive CD8^+^ T cells were all at least 2-fold greater in ND mice than in HFD mice (2.4-, 2.0-, and 2.9-fold, respectively). Overall, significant CD8^+^ T cell activation was observed in ND but not HFD mice that received the SARS-CoV-2 BNT162b2 mRNA vaccine, indicating an impaired antigen-specific CD8^+^ T cell Spike response in HFD mice.

**Figure 3.**
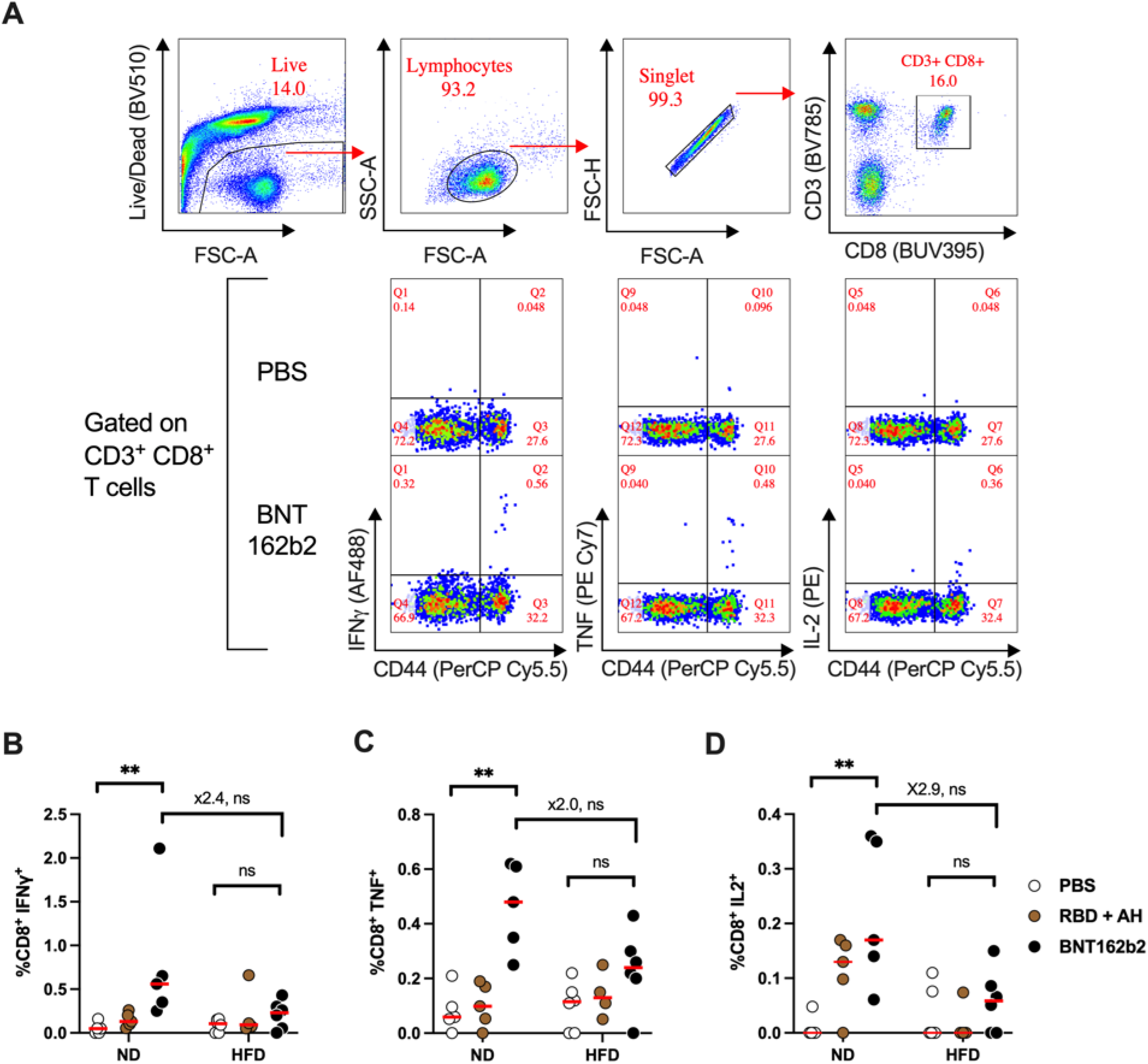
SARS-CoV-2 mRNA vaccine enhances CD8^+^ T cell responses in ND but not HFD mice. Male C57BL/6J mice fed with high-fat diet (HFD) or normal diet (ND) were immunized as in Figure 2. Splenocytes were collected 4 weeks after the final immunization and stimulated with a SARS-CoV-2 spike RBD peptide pool. (**A**) Representative flow data. (**B-D**) Expression of intracellular IFN_γ_ (**B**), TNF (**C**), and IL-2 (**D**) was assessed by flow cytometry in CD8^+^ T cells. **N** = 4-6 per group. Bars represent median. Dots represent individual values. Data were analyzed by the Kruskal-Wallis Test adjusted for multiple comparisons. Fold difference between BNT162b2-immunized ND and HFD mice are shown. ** *P* < 0.01.

### SARS-CoV-2 mRNA vaccine protects ND mice but not HFD mice from lung infection

To assess vaccine efficacy, we challenged immunized mice with live SARS-CoV-2. Eight weeks after the final immunization, mice were challenged intranasally with 10^3^ PFU of mouse-adapted MA10 SARS-CoV-2^27^, as indicated in **Fig. 2A**. Mice were weighed before infection, and the HFD mice remained significantly heavier, with a mean weight of 52.7 g versus a mean of 34.0 g among ND mice (*P* < 0.0001). Two days after infection, mice were euthanized, and lungs were harvested for analysis. Minor inflammation was seen in all lung samples, though differences between groups were undetectable in line with the early timepoint following infection (**fig. S3**).

To determine protective efficacy, host lung inflammatory responses were evaluated by assessing gene expression of cytokines, chemokines, and IFN-stimulated genes (ISGs) associated with SARS-CoV-2 severity including *Ifit2, Cxcl10, Csf2, Il6, Ccl2* and *Cxcl1*^28-32^. PBS-injected HFD and ND mice both demonstrated high inflammatory responses across multiple genes (**Fig. 4A**). Of note, BNT162b2-immunized ND mice demonstrated significant lower gene expression relative to PBS group, while there was no significant difference between HFD mice that received PBS or BNT162b2 (**Fig. 4A**). To further determine protective efficacy, lung viral titers were analyzed (**Fig. 4B**). Naive, PBS-injected mice demonstrated robust viral loads in both HFD and ND mice (geometric means: 1.58 × 10^8^ and 6.20 × 10^9^, respectively). In line with the low immunogenicity data, both HFD and ND mice vaccinated with alum-adjuvanted RBD were not protected from lung infection (geometric means: 2.61 × 10^7^ and 3.80 × 10^8^ in HFD and ND mice respectively, **Fig. 4B**). Similar to the pattern of expression observed for inflammatory genes, ND mice that received BNT162b2 exhibited a significant decrease in lung viral titer relative to the PBS group (*P* = 0.004), while there was no significant difference between HFD mice that received PBS or BNT162b2 (*P* = 0.535, **Fig. 4B**). Compared to HFD, ND mice that received BNT162b2 demonstrated a 262-fold reduction in lung viral titers (geometric means: 9.25 × 10^6^ and 3.53 × 10^4^ in HFD and ND mice, respectively), though this difference was not statistically significant (*P* = 0.175, **Fig. 4B**). Lastly, to define immune correlates of protection, we assessed correlations between neutralizing titers and challenge study readouts. Neutralizing titers demonstrated strong inverse correlations with *Ifit2* gene expressions (*r* = −0.8166, *P* < 0.0001) and lung viral loads (*r* = −0.6640, *P* = 0.0002) (**Fig. 4C**). Although one HFD mouse with high neutralizing titer demonstrated high lung viral titer, the mouse was protected from lung inflammation with suppressed *Ifit2* expression (arrow, **Fig 3C**). Overall, these results demonstrate that HFD mice are not protected from infection with SARS-CoV-2 following immunization with BNT162b2, while ND mice are significantly protected against viral lung infection after receiving BNT162b2.

**Figure 4:**
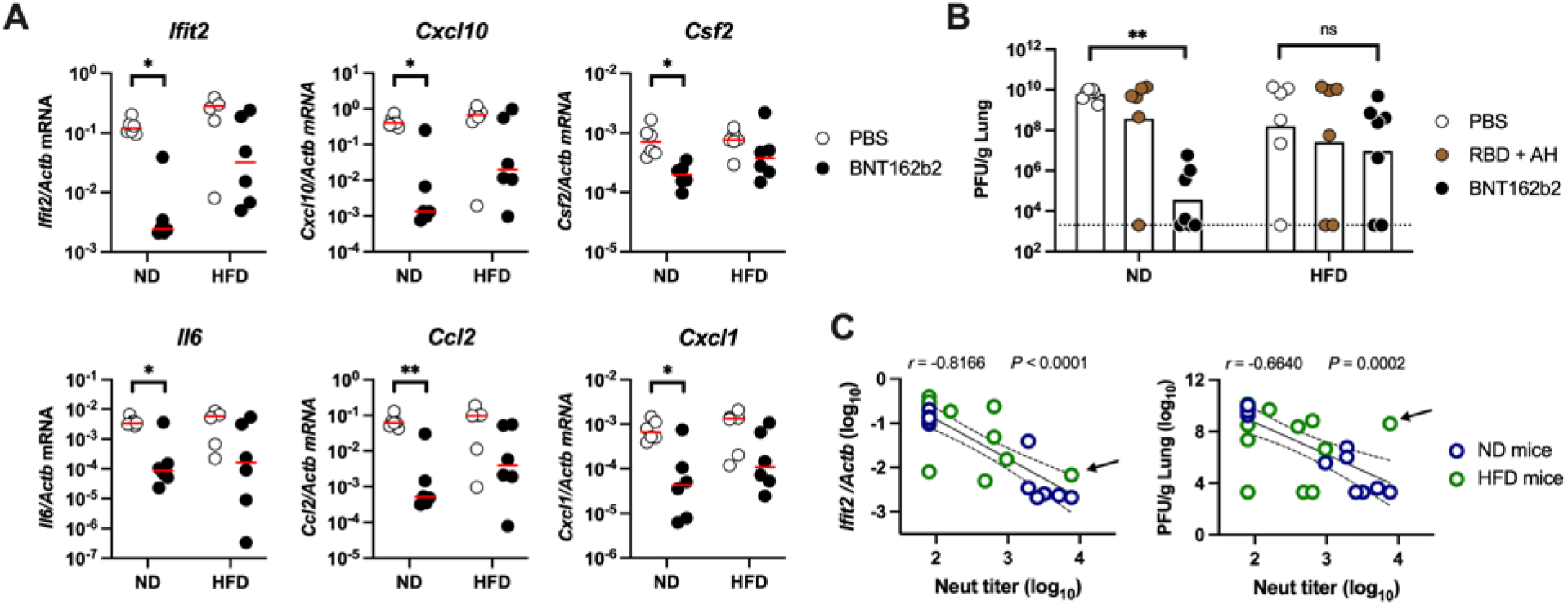
SARS-CoV-2 mRNA vaccine protects ND mice but not HFD mice. Male C57BL/6J mice fed high-fat diet (HFD) or normal diet (ND) were immunized as in Figure 2. Six weeks after immunization, mice were challenged with 10^3^ PFU of MA10 SARS-CoV-2. Mice were euthanized 2 days after infection and lungs were harvested for analysis. (**A, B**) To assess protective efficacy, lung homogenates were analyzed for gene expression profiles of *Ifit2, Cxcl10, Csf2, Il6, Ccl2* and *Cxcl1* shown as relative expression compared to *Actb* (**A**) and viral titers (**B**). Bars represent medians (**A**) or geometric means (**B**). Dashed lines represent lower limits of detection. n = 6–7 per group. Data were analyzed by the Mann-Whitney or Kruskal–Wallis test corrected for multiple comparisons. \**P* < 0.05, \*\**P* < 0.01. (**C**) Correlations between neutralizing titers and gene expressions of *Ifit2* over *Actb* and lung viral loads are shown. Circles represent individual mice that received PBS or BNT162b2, and colors indicate ND or HFD mice. Solid and dashed lines respectively indicate linear regression and 95% confidence interval. Correlations were assessed by two-sided Pearson tests. Arrow represents one HFD mouse with high neutralizing titer showing high lung viral titer but suppressed *Ifit2* expression.

## DISCUSSION

Overall, we have shown for the first time that HFD-induced insulin resistance and obesity impair SARS-CoV-2 mRNA vaccine humoral and cellular immunogenicity, providing causal evidence to support observations in human patients and establishing a model for studying the relationship between metabolic diseases and SARS-CoV-2 vaccine responses. T2DM and obesity are known risk factors for severe COVID-19 and have been correlated with reduced responses to mRNA vaccines against SARS-CoV-2^7-9, 33^. However, a causal link between these conditions and impaired vaccine responses has not yet been established. Using DIO mouse models, previous studies have recapitulated pathological findings of MERS-CoV and SARS-CoV-2 observed in humans, establishing this model as a viable option for studying the relationship between metabolic disease and SARS-CoV-2 vaccine response^19-21^. Here, we developed a mouse model of metabolic disease by feeding mice a HFD, which led to obesity, hyperinsulinemia, and glucose intolerance. We then immunized mice with SARS-CoV-2 mRNA BNT162b2 vaccine, an alum-adjuvanted RBD subunit vaccine, or a PBS control, after which we assessed Ab and T cell responses and protection from viral challenge. We found that HFD mice exhibited an overall reduction in BNT162b2 response relative to ND mice, marked by a reduction in neutralizing Abs. We also observed significantly enhanced RBD-specific CD8^+^ T cell induction only in ND mice but not in HFD mice. Furthermore, BNT162b2-vaccinated HFD mice exhibited a lack of protection from live viral challenge relative to PBS-injected HFD mice, while BNT162b2-vaccinated ND mice demonstrated protection relative to the PBS-injected ND mice.

We observed that in HFD mice, BNT162b2 vaccine demonstrated a consistent and cumulative pattern of reduced immunogenicity across multiple measures, including binding and neutralizing Ab titers and CD8^+^ T cell activation. Interestingly, while antigen-specific IgG1 titers were comparable among HFD and ND mice, the induction of IgG2c Abs was substantially reduced in HFD mice relative to ND mice. We also observed that, unlike ND mice, HFD mice failed to mount significant mRNA vaccine-induced CD8^+^ T cell activation and expression of Th1-associated cytokines including IFNγ, associated with favorable disease outcomes^26, 34^. As IFNγ promotes isotype switching toward IgG2c *in vivo*^*35*^, these two observations are likely linked. Our study is consistent with impaired CD8^+^ T cell responses following influenza vaccination and natural infection in HFD mice^11, 13, 16^. Furthermore, susceptibility to infection in HFD mice may in part be due to impaired generation of IgG2c Ab subclass, associated with greater effector functions (e.g., induction of phagocytosis, complement fixation) likely important for host defense against infection^36^. Overall, our data demonstrate that HFD obese and diabetic mice have distinct immunity with impaired generation of neutralizing Abs and IFNγ-driven type 1 immunity.

To evaluate whether immunogenicity data translate into protection, we performed a live challenge study. Here, we observed a significant reduction in lung inflammatory responses and lung viral titers relative to PBS-injected mice at day 2 post-infection among ND mice but not in HFD mice. As expected, and in line with immunogenicity data, this result demonstrates that BNT162b2-immunized HFD mice are not protected from challenge, while ND mice are mostly protected. Notably, we did not observe worse disease outcomes (i.e., high viral titers and lung inflammatory responses) in naive, PBS-injected HFD mice compared to ND mice, despite prior observations of more severe disease in HFD animals^19^. However, the shorter duration of follow-up post challenge, which was chosen to maximize observable differences in lung viral titers, likely accounts for these discrepancies.

By establishing a causal connection between metabolic disease and vaccine efficacy, our study lays the groundwork for future inquiries into the mechanisms behind diminished vaccine responses. T2DM and obesity are characterized by chronic low-grade inflammation, also known as ‘metaflammation’^37-41^, sharing features with ‘inflammaging’, the chronic, sterile, low-grade, inflammatory state that characterizes aging^41, 42^. Metaflammation and inflammaging both contribute to the key pathogenesis of metabolomic- or age-related diseases–however, the association and its mechanism on vaccine immunogenicity are not fully elucidated. Senescent cells in older adults provoked CCR2 positive monocyte-dependent inflammation and diminished T cell responses to viruses via secretion of prostaglandin E_2_^43^. Interestingly, a short-term inhibition of inflammatory responses boosted adaptive immunity in aged mice^43^. Additionally, T2DM-induced insulin resistance in humans has been linked with impaired ability for CD14^+^ monocytes to differentiate into dendritic cells (DCs), which then also show reduced classical DC maturation and antigen presenting function^44^. In light of these published studies, our overall findings suggest that hyper-inflammatory states associated with obesity and type 2 DM likely mediated the observed deficits in vaccine response among HFD mice. Future studies should elucidate the mechanistic connections between metabolic disease and vaccine immunogenicity, which could enable implementation of targeted strategies to overcome deficits in vaccine response in vulnerable populations with distinct immunity.

While SARS-CoV-2 vaccines tailored for those with obesity or T2DM do not yet exist, strategies to develop precision vaccines for specific age populations have been investigated. In line with our approach to modeling metabolic disease, age-specific murine models have demonstrated reduced immunogenicity, higher mortality and morbidity, and greater waning immunity in aged mice, comparable to the observations in older adult humans^45-47^. A booster of mRNA vaccine provided sterilizing immunity against Omicron-induced lung infection in aged 21-month-old mice, while younger mice are protected without a booster, indicating the importance of age-specific vaccine regimens^46^. Through the development of an appropriate adjuvant for a SARS-CoV-2 protein-based vaccine, greater protection has been observed in aged mice despite age-related declines in immunity^47^. Based on this precedent, we hypothesize that similar approaches could help overcome metabolic disease-associated deficits in vaccine response. Of note, adjuvants can not only enhance vaccinal immunity but also shape the polarization of the immune response^47, 48^. Defining optimal adjuvant formulation could therefore be a promising approach to overcome the reduced Th1 polarization observed among diabetic obese mice in this study. In combination with further studies elucidating the mechanisms of impaired vaccine responses among those with metabolic disease, an adjuvant approach therefore represents a promising future direction toward effective vaccines tailored to those with T2DM and obesity^49^.

Our study has several major strengths, including (a) the comprehensive assessment of a causal connection between metabolic disease and reduced BNT162b2 immunogenicity among neutralizing Abs, IgG2c subclasses, and cytotoxic T cells, and (b) evaluation of protective efficacy from live SARS-CoV-2 challenge. Despite these strengths, we recognize several limitations in the current study, including that (a) only male mice were used due to the increased severity of obesity and insulin resistance in male C57BL/6J mice relative to females^50, 51^, (b) only one mouse model was used, establishing the need for future translational research in additional animal models and humans, (c) the overall magnitude of antigen-specific T cell responses were low even after mRNA vaccination due to the use of RBD-specific peptide pool instead of full spike-peptide pool, and (d) although we showed the association with HFD mice and an impairment of type 1 immunity, we were not able to demonstrate the contribution for protection as we had to euthanize mice and collect splenocytes to assess T cell response. Nevertheless, the implications of metabolic disease on BNT162b2 immunogenicity are clear, laying the groundwork for further study into the mechanisms of impaired immune responses, especially focused on a) insulin resistance and b) methods for overcoming these phenomena both in animal models and eventually in the clinic. In parallel to this precision vaccinology approach, public health initiatives that promote physical exercise and a health body weight, both know to help curtail insulin resistance, should be adopted.

Overall, this study aimed to analyze the effects of obesity and insulin resistance on immunogenicity and protective efficacy following immunization with SARS-CoV-2 BNT162b2 mRNA vaccine. We demonstrated that HFD-induced obesity and insulin resistance led to reduced humoral and cellular immunogenicity of the BNT162b2 vaccine. Furthermore, a weakened protective efficacy was shown in HFD mice post BNT162b2 immunization. These observations establish the need to develop precision vaccines against SARS-CoV-2 and other pathogens tailored for those with obesity and DM to overcome impaired immune responses in groups already at high risk of severe infections^52^.

## MATERIALS AND METHODS

### Study design

This study aimed to assess the effects of diet-induced obesity and insulin resistance on BNT162b2 mRNA SARS-CoV-2 vaccine immunogenicity and protection in pre-clinical mouse models. To this end, we used longitudinal mouse *in vivo* models fed either a high-fat or control diet to dissect the effects of the high-fat diet and associated phenotypes on vaccine immunogenicity and infection protection. Sample size was chosen empirically based on the results of previous studies and practical limitations such as vivarium capacity. The *in vivo* arm of the study was completed over a single 9-month period, with animal husbandry and associated procedures completed by the same staff throughout. Mouse experiments aimed to include a total of 12–13 mice per group. Mice were randomly assigned to different treatment groups. No data outliers were excluded.

### Animals

Male, 14–15-week-old C57BL/6J mice fed on a high-fat or control diet beginning at age 6 weeks were purchased from Jackson Laboratory. Mice were housed under specific pathogen-free conditions at Boston Children’s Hospital, and all the procedures were approved under the Institutional Animal Care and Use Committee (IACUC) and operated under the supervision of the Department of Animal Resources at Children’s Hospital (ARCH) (Protocol number 00001573). Mice were fed either a high-fat diet containing 60% kcal from fat (D12492i, Research Diets) or an ingredient-matched control diet containing 10% kcal from fat (D12450Ji, Research Diets) from age 6 weeks until the end of the study. At the University of Maryland School of Medicine, mice were housed in a biosafety level 3 (BSL3) facility for all SARS-CoV-2 infections with all the procedures approved under the IACUC (Protocol number #1120004) to MBF.

### Fasting Insulin ELISA

Mice were transferred to clean cages without food and fasted for 6 hours. Blood was collected via retro-orbital bleed and serum was isolated by centrifugation at 1500 g for 7.5 minutes. Serum insulin was measured by ELISA according to the manufacturer’s wide-range detection protocol (Crystal Chem).

### Intraperitoneal glucose tolerance test

An intraperitoneal glucose tolerance test was performed by adapting an existing protocol ^22^. Briefly, mice were transferred to clean cages without food, weighed, and fasted for 6 hours. Following fasting, mice were restrained, blood was drawn from the tail vein using a 30-gauge lancet, and baseline blood glucose was measured using a OneTouch Verio Flex meter (LifeScan). A 20% sterile dextrose solution (ICU Medical) was administered via intraperitoneal injection at a final concentration of 2 g dextrose/kg. Blood glucose was measured at 30, 75, and 120 minutes after injection. The resulting values were recorded, and any measurements over the meter’s upper limit of detection (600 mg/dL) were assigned a value of 600 mg/dL.

### SARS-CoV-2 wildtype spike and RBD expression and purification

Full length SARS-CoV-2 Wuhan-Hu-1 spike glycoprotein (M1-Q1208, GenBank MN90894) and RBD constructs (amino acid residues R319-K529, GenBank MN975262.1), both with an HRV3C protease cleavage site, a TwinStrepTag and an 8XHisTag at C-terminus were obtained from Barney S. Graham (NIH Vaccine Research Center) and Aaron G. Schmidt (Ragon Institute), respectively. These mammalian expression vectors were used to transfect Expi293F suspension cells (Thermo Fisher) using polyethylenimine (Polysciences). Transfected cells were allowed to grow in 37°C, 8% CO_2_ for an additional 5 days before harvesting for purification. Protein was purified in a PBS buffer (pH 7.4) from filtered supernatants by using either StrepTactin resin (IBA) or Cobalt-TALON resin (Takara). Affinity tags were cleaved off from eluted protein samples by HRV 3C protease, and tag removed proteins were further purified by size-exclusion chromatography using a Superose 6 10/300 column (Cytiva) for full length Spike and a Superdex 75 10/300 Increase column (Cytiva) for RBD domain in a PBS buffer (pH 7.4).

### Adjuvants and immunization

BNT162b2 suspension (100 µg/mL) was diluted 1:5 in PBS, and 1 µg of mRNA was injected. Mice in the RBD + aluminum hydroxide condition received 10 µg of recombinant monomeric SARS-CoV-2 RBD protein formulated with 100 µg of Alhydrogel adjuvant 2% (Invivogen). Mice in the PBS vaccination group received phosphate-buffered saline (PBS) alone. BNT162b2 spike mRNA vaccine (Pfizer-BioNTech) was obtained as otherwise-to-be-discarded residual volumes in used vials from the Boston Children’s Hospital vaccine clinic and was used within 6 hours from the time of reconstitution. Injections (50 µL) were administered intramuscularly in the caudal thigh on days 0 and 14. Blood samples were collected 2 weeks post-immunization.

### Antibody ELISA

RBD- and spike protein-specific Ab concentrations were quantified in serum samples by ELISA using a previously described protocol ^53^. Briefly, high-binding flat-bottom 96-well plates (Corning) were coated with 50 ng per well RBD or 25 ng per well spike protein and incubated overnight at 4 °C. Plates were washed with 0.05% Tween 20 PBS and blocked with 1% bovine serum albumin (BSA) in PBS for 1 hour at room temperature. Serum samples were serially diluted 4-fold from 1:100 up to 1:1.05 × 10^8^ and then incubated for 2 hours at room temperature. Plates were washed three times and incubated for 1 hour at room temperature with horseradish peroxidase (HRP)-conjugated anti-mouse IgG, IgG1, IgG2a, or IgG2c (Southern Biotech). Plates were washed five times and developed with tetramethylbenzidine (1-Step Ultra TMB-ELISA Substrate Solution, Thermo Fisher Scientific, for the RBD ELISA, and BD OptEIA Substrate Solution, BD Biosciences, for the spike ELISA) for 5 minutes, then stopped with 2 N H_2_SO_4_. Optical densities (ODs) were read at 450 nm with a SpectraMax iD3 microplate reader (Molecular Devices). End-point titers were calculated as the dilution that emitted an optical density exceeding a 3× background. An arbitrary value of 50 was assigned to the samples with OD values below the limit of detection for which it was not possible to interpolate the titer.

### hACE2-RBD inhibition assay

The hACE2-RBD inhibition assay modified a previously existing protocol ^47, 54^. Briefly, high-binding flat-bottom 96-well plates (Corning) were coated with 100 ng per well recombinant human ACE2 (hACE2) (Sigma-Aldrich) in PBS, incubated overnight at 4°C, washed three times with 0.05% Tween 20 PBS, and blocked with 1% BSA PBS for 1 hour at room temperature. Each serum sample was diluted 1:80, pre-incubated with 3 ng of RBD-Fc in 1% BSA PBS for 1 hour at room temperature, and then transferred to the hACE2-coated plate. RBD-Fc without pre-incubation with serum samples was added as a positive control, and 1% BSA PBS without serum pre-incubation was added as a negative control. Plates were then washed three times and incubated with HRP-conjugated anti-human IgG Fc (Southern Biotech) for 1 hour at room temperature. Plates were washed five times and developed with tetramethylbenzidine (BD OptEIA Substrate Solution, BD Biosciences) for 5 min, then stopped with 2 N H_2_SO_4_. The optical density was read at 450 nm with a SpectraMax iD3 microplate reader (Molecular Devices). Percentage inhibition of RBD binding to hACE2 was calculated with the following formula: Inhibition (%) = [1 – (Sample OD value – Negative Control OD value)/(Positive Control OD value – Negative Control OD value)] × 100.

### SARS-CoV-2 neutralization titer determination

All serum samples were heat-inactivated at 56°C for 30 min to deactivate complement and allowed to equilibrate to RT prior to processing for neutralization titer. Samples were diluted in duplicate to an initial dilution of 1:40 followed by 1:2 serial dilutions, resulting in a 12-dilution series with each well containing 60 µl. All dilutions employed DMEM (Quality Biological), supplemented with 10% (v/v) fetal bovine serum (heat-inactivated, Gibco), 1% (v/v) penicillin/streptomycin (Gemini Bio-products) and 1% (v/v) L-glutamine (2 mM final concentration, Gibco). Dilution plates were then transported into the BSL-3 laboratory and 60 µl of diluted SARS-CoV-2 (WA-1, courtesy of Dr. Natalie Thornburg/CDC) inoculum was added to each well to result in a multiplicity of infection (MOI) of 0.01, or 100 pfu/well, upon transfer to titering plates and an initial serum dilution with virus added of 1:80. A non-treated, virus-only control and mock infection control were included on every plate. The sample/virus mixture was then incubated at 37°C (5.0% CO_2_) for 1 hour before transferring 100 µl to 96-well titer plates with 1e4 VeroTMPRSS2 cells. Titer plates were incubated at 37°C (5.0% CO_2_) for 72 hours, followed by cytopathic effect (CPE) determination for each well in the plate. The first sample dilution to show CPE was reported as the minimum sample dilution required to neutralize >99% of the concentration of SARS-CoV-2 tested (NT_99_).

### Splenocyte restimulation, intracellular cytokine staining and flow cytometry

Mouse spleens were mechanically dissociated and filtered through a 70 µm cell strainer. After centrifugation, cells were treated with 1 mL ammonium-chloride-potassium lysis buffer for 2 minutes at RT. Cells were washed and plated in a 96-well U-bottom plate (2 × 10^6^/well) and rested overnight at 37 ºC in RPMI 1640 supplemented with 10% heat-inactivated FBS, penicillin (100 U/ml), streptomycin (100 mg/ml), 2-mercaptoethanol (55 mM), non-essential amino acids (60 mM), HEPES (11 mM), and L-Glutamine (800 mM) (all Gibco). Next day, SARS-CoV-2 RBD peptide pools (PM-WCPV-S-RBD-1, JPT) were added at 0.6 nmol/ml in the presence of anti-mouse CD28/49d (1 μg/mL, BD) and brefeldin A (5 μg/ml, BioLegend). After a 6-hour stimulation, cells were washed twice and treated with Mouse Fc Block (BD) according to the manufacturer’s instructions. Cells were washed and stained with Aqua Live/Dead stain (Life Technologies, 1:500) for 15 minutes at RT. Following two additional washes, cells were incubated with the following Abs for 30 minutes at 4°C: anti-mouse CD44 [IM7, PerCP-Cy5.5, BioLegend #103032, 1:160], anti-mouse CD3 [17A2, Brilliant Violet 785, BioLegend #100232, 1:40], anti-mouse CD4 [RM4-5, APC/Fire 750, BioLegend 100568, 1:160] and anti-mouse CD8 [53-6.7, Brilliant UltraViolet 395, BD #563786, 1:80]. Cells were then fixed and permeabilized by using the BD Cytofix/Cytoperm kit according to the manufacturer’s instructions and were subjected to intracellular staining (30 minutes at 4 °C) using the following Abs: anti-mouse IFNγ [XMG1.2, Alexa Fluor 488, BioLegend #505813, 1:160], anti-mouse TNF [MP6-XT22, PE Cy7, BioLegend # 506324, 1:160], anti-mouse IL-2 [JES6-5H4, PE, BioLegend # 503808, 1:40]. Finally, cells were fixed in 1% paraformaldehyde (Electron Microscopy Sciences) for 20 minutes at 4 ºC and stored in PBS at 4 ºC until acquisition. Samples were analyzed on an LSR Fortessa (BD) flow cytometer and FlowJo v10.8.1 (FlowJo LLC).

### SARS-CoV-2 mouse challenge study

Mice were anesthetized by intraperitoneal injection of 50 μL of a mix of xylazine (0.38 mg/mouse) and ketamine (1.3 mg/mouse) diluted in PBS. Mice were then intranasally inoculated with 1 × 10^3^ PFU of mouse-adapted SARS-CoV-2 (MA10, courtesy of Dr. Ralph Baric (UNC)) in 50 μL divided between nares ^27^. Challenged mice were weighed on the day of infection and daily for up to 2 days post-infection. At 2 days post-infection, mice were euthanized, and lungs were harvested to determine virus titer by a plaque assay and prepared for histological staining and RNA extraction.

### SARS-CoV-2 plaque assay

The day prior to infection, 2.5e5 VeroTMPRSS2 cells were seeded per well in a 12-well plate in 1mL of VeroTMPRSS2 media. Tissue samples were thawed and homogenized with 1mm beads in an Omni Bead ruptor (Omni International Inc., Kennesaw, GA) and then spun down at 21,000 g for 2 minutes. A 6-point dilution curve was prepared by serial diluting 25 μL of sample 1:10 in 225 μL DMEM. 200 μL of each dilution was then added to the cells and the plates were rocked every 15 minutes for 1 hour at 37°C. After 1 hr, 2 mL of a semi-solid agarose overlay was added to each well (DMEM, 4% FBS, 0.06% UltraPure agarose (Invitrogen, Carlsbad, CA). After 48 hours at 37°C and 5% CO_2_, plates were fixed in 2% PFA for 20 minutes, stained with 0.5 mL of 0.05% Crystal Violet and 20% EtOH, and washed 2x with H_2_O prior to counting of plaques. The titer was then calculated. For tissue homogenates, this titer was multiplied by 40 based on the average tissue sample weight being 25 mg.

### Gene expression analysis by qPCR

RNA was isolated from TRI Reagent samples using phenol-chloroform extraction or column-based extraction systems (Direct-zol RNA Miniprep, Zymo Research) according to the manufacturer’s protocol. RNA concentration and purity (260/280 and 260/230 ratios) were measured by NanoDrop (Thermo Fisher Scientific). Samples with an A260/A280 ratio of <1.8 were excluded for further analysis. cDNA was prepared from purified RNA with RT2 First Strand Kit, per the manufacturer’s instructions (Qiagen). cDNA was quantified by qPCR on a 7300 real-time PCR system (Applied Biosystems) using pre-designed SYBR Green Primers (QIAGEN) specific for *Ifit2* (PPM05993A), *Cxcl10* (PPM02978A), *Csf2* (PPM02990A), *Il6* (PPM03015A), *Ccl2* (PPM03151A), *Cxcl1* (PPM03058A), and *Actb* (PPM02945A).

### Histopathology analysis

Slides were prepared as 5 μm sections and stained with hematoxylin and eosin. A pathologist was blinded to information identifying the treatment groups and fields were examined by light microscopy.

### Statistical analysis

Statistical analyses employed Prism v9.4.0 (GraphPad Software). *P* values < 0.05 were considered significant. Normally distributed data were analyzed by t-test or one- or two-way analyses of variance (ANOVAs). To achieve normal distribution, some datasets were analyzed after Log-transformation as indicated in the figure legends. Non-normally distributed data were analyzed by Mann-Whitney U-test or Kruskal-Wallis test. *P* values were corrected for multiple comparisons.

## Abbreviations

T2DM: Type 2 diabetes mellitus
DIO: diet-induced obesity
Ab: antibody
HFD: high-fat diet
RBD: receptor-binding domain
ND: normal diet
IPGTT: intraperitoneal glucose tolerance test
IFN: interferon
ISGs: IFN-stimulated genes
DC: dendritic cell

## LIST OF SUPPLEMENTARY MATERIALS

Figures S1 to S3

Table S1

## ACKNOWLEDGEMENTS

We thank the members of the BCH *Precision Vaccines Program* (PVP) for helpful discussions as well as Kevin Churchwell, Gary Fleisher, David Williams, and August Cervini for their support of the PVP. We thank Ralph Baric for providing the SARS-CoV-2/MA10 virus. We thank B. S. Graham (NIH Vaccine Research Center) for providing the plasmid for prefusion stabilized SARS-CoV-2 spike trimer and Aaron G. Schmidt for providing the spike RBD constructs used for protein expression. We thank the pharmacists of BCH for their efforts to maximize the use of SARS-CoV-2 vaccines by saving leftover or to-be-discarded overfill from BNT162b2 vaccine vials. E.N. is a JSPS Overseas Research Fellow and a joint Society for Pediatric Research and Japanese Pediatric Society Scholar. D.J.D. thanks Siobhan McHugh, Geneva Boyer, Lucy Conetta, and the staff of Lucy’s Daycare, the staff of YMCA of Greater Boston, Bridging Independent Living Together (BILT), Inc., and the Boston Public Schools for childcare and educational support during the COVID-19 pandemic. The graphics in **Fig.1A** and **2A** were created with BioRender.

---

**Supplementary Table 1.**
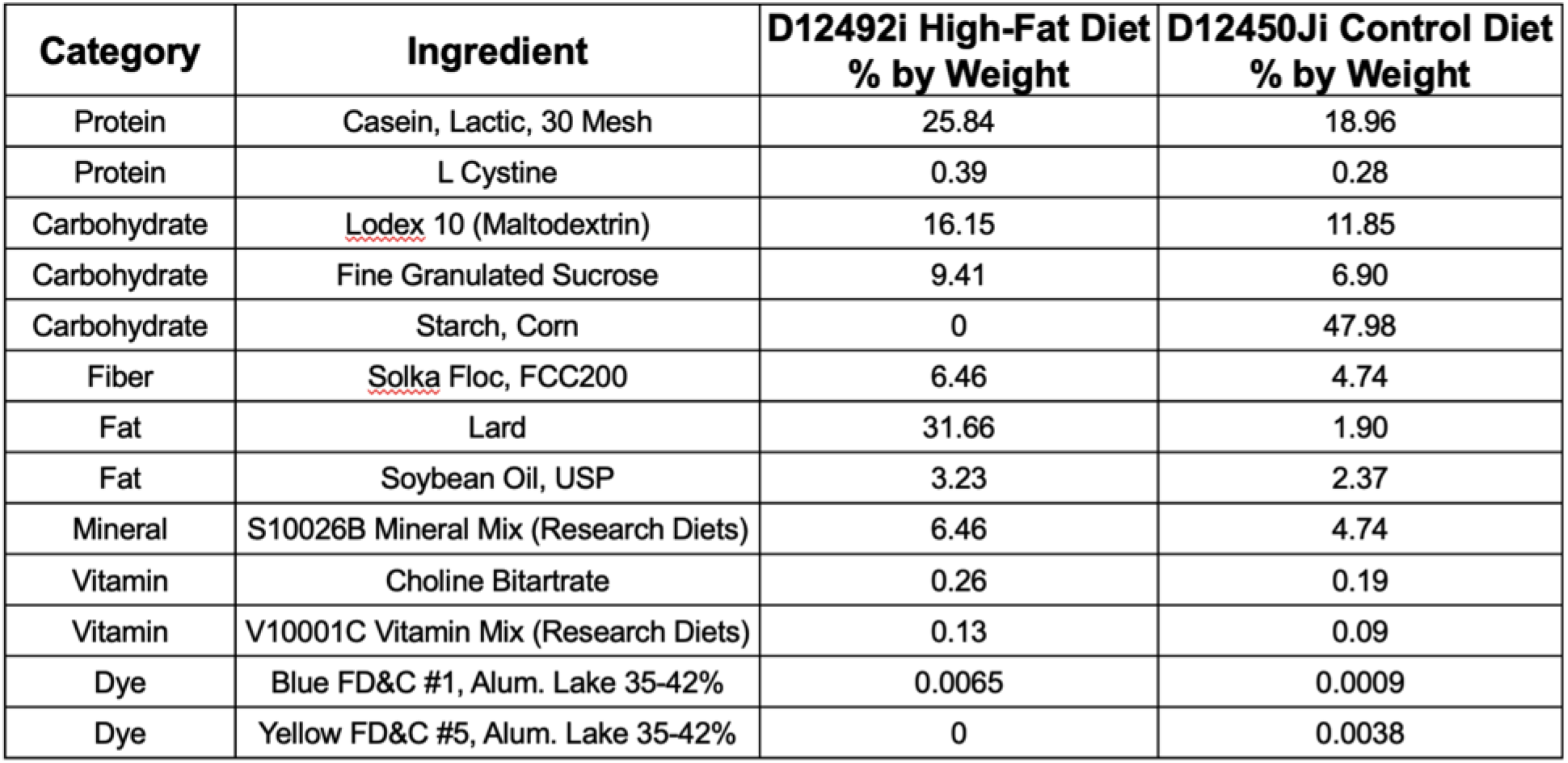
Composition of high-fat and control diets. High-fat diet D12492i (Research Diets) comprised of 60% kcal from fat and control diet D12450Ji (Research Diets) comprised of 10% kcal from fat were fed to male C57BL/6J mice.

**Supplementary Figure 1.**
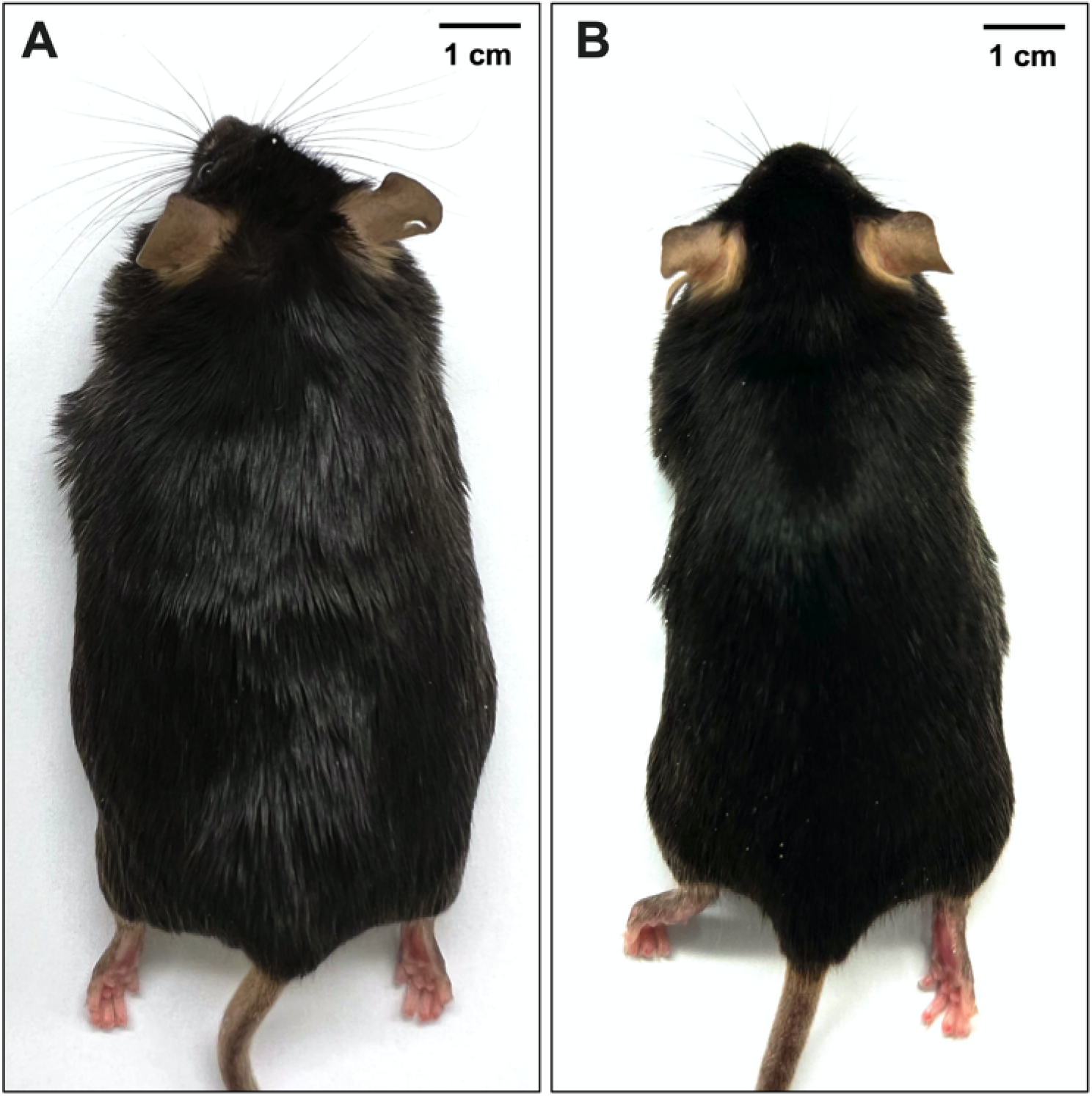
A high-fat diet induces obesity in male C57BL/6J mice. Representative images of (A) diet-induced obese and (B) control male C57BL/6J mice at age 30 weeks.

**Supplementary Figure 2.**
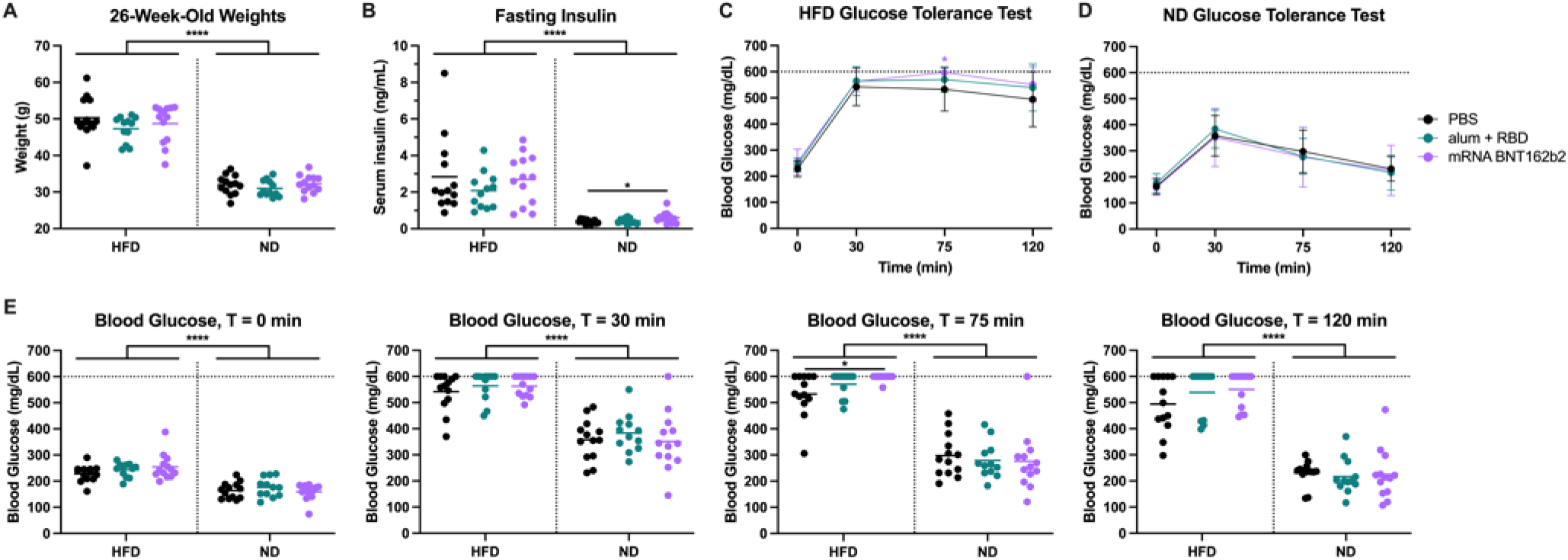
Obesity, hyperglycemia, and poor glucose tolerance phenotypes are comparable across vaccine treatment groups within each strain. (A) Mice were weighed during the week of the first immunization. Weights are grouped by vaccine condition and mouse diet (HDF or ND). (B) Serum insulin levels were measured after a 6-hour fast. (C–E) Glucose tolerance was assessed by measuring blood glucose at time points 0, 30, 75, and 120 min following a 6-hour fast and intraperitoneal injection of 2 g/kg dextrose. Blood glucose values above the glucometer’s 600 mg/dL upper limit of detection were assigned a value of 600 mg/dL. Longitudinal graphs display mean and SD of HFD (C) and ND (D) strains. Bars represent means in all dotplots. Horizontal dotted lines represent upper limits of detection. n = 13 per group in the PBS and mRNA BNT 162b2 groups and n = 12 in the alum + RBD group across both strains. Significance within each strain was assessed by Kruskal-Wallis test with post-hoc Dunn’s multiple comparisons test. Comparisons between HFD and ND mice were assessed by unpaired t-test (A, B) or Mann-Whitney U tests (E). * *P* < 0.05 and **** *P* < 0.0001.

**Supplementary Figure 3.**
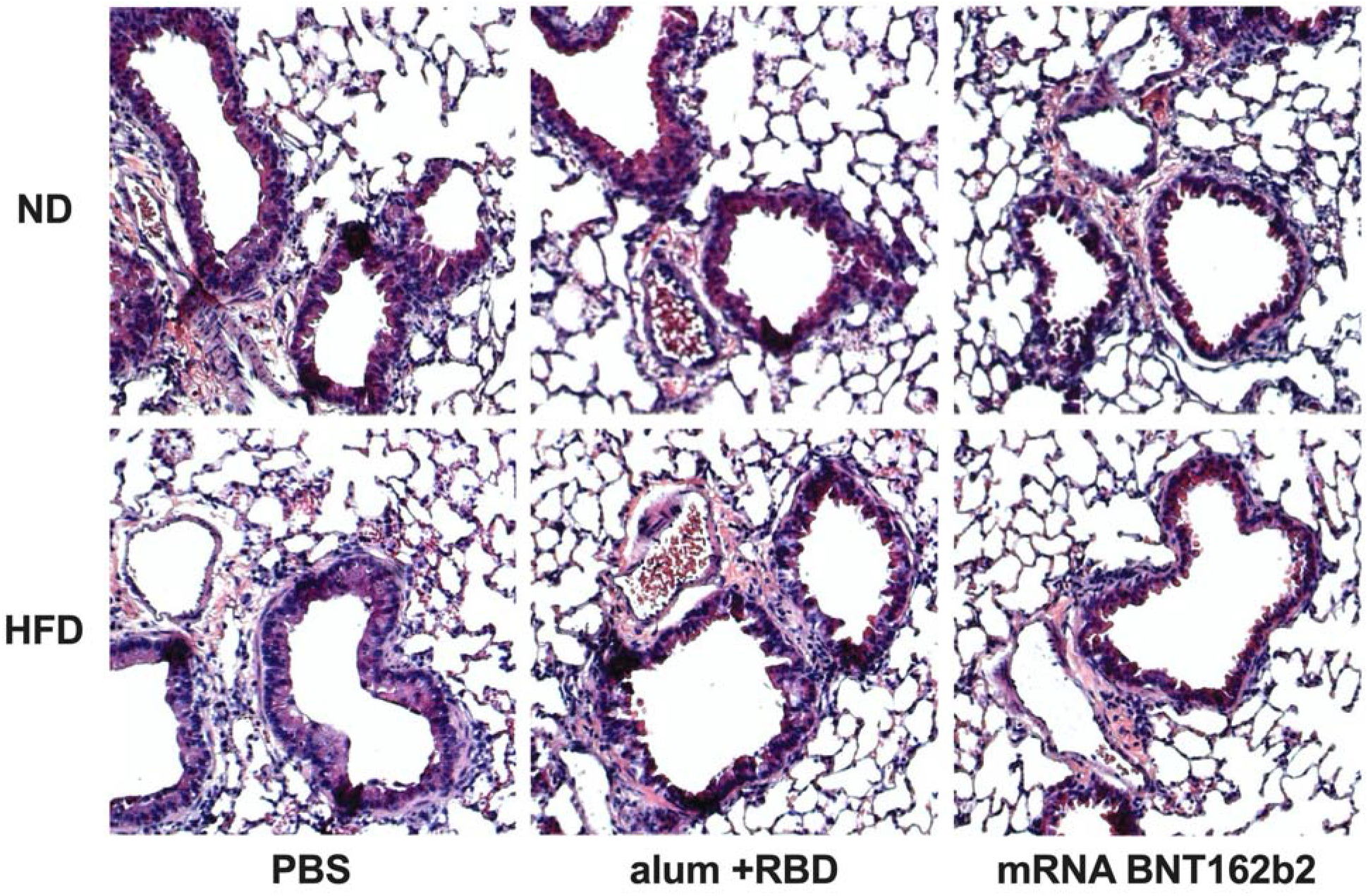
Lung histopathology following live SARS-CoV-2 MA10 challenge. Lung tissue was harvested at 2 days post challenge, fixed, sectioned, and stained using hematoxylin and eosin. Representative images are shown. N = 6–7 per group.

